# Mitochondrial diversity in the ribbed mussel, *Geukensia demissa*, relative to high marsh plant diversity at the southern edge of the distribution

**DOI:** 10.1101/2022.01.16.476515

**Authors:** Ivy G. Francis, John P. Wares

## Abstract

Genomic diversity and relatedness among sample sites are often used to explore landscape-level processes of how and where organisms are limited in movement. In many cases, these patterns of diversity and relatedness can be useful for understanding larger ecological patterns. A prior study has suggested that larval input – inferred from landscape genomic data – of the ribbed mussel *Geukensia demissa*, a species with important roles in stabilizing salt marsh ecosystems, could be indicative of longer-term recruitment patterns of high marsh plant species. Here we use new observations of mitochondrial diversity in the same region but with more sites sampled to show that this prior study was wrong in suggesting that relationship. The same mitochondrial data are useful for monitoring cryptic patterns of climate response in these mussels relative to a subtropical congener.

---

The diversity contained within populations may be represented by individual traits, behaviors, phenotypes, or genomes (Bolnick et al. 2003). A key transition in our understanding of this diversity has taken place in the past two decades - we have abundant data that species are often not spatially homogeneous, and the collective patterns of diversity tell us about processes and forecasts for how a species will respond as the environment changes (Hughes and Stachowicz 2004). Understanding how diversity in species shapes or interacts with the environment around it is of particular importance when the species plays an important foundational or keystone role in the community. Such studies have given insight into declining sexual recruitment in some coral communities (Baums et al 2006), fitness-based transitions within species distributions (Wares et al 2021), and the best practices of management and recovery for economically important species like oysters (Downey-Wall et al 2020).

The ribbed mussel, *Geukensia demissa*, has been recognized as playing such a fundamental role in Atlantic salt marsh ecosystems. These mussels recruit to salt marshes and attach themselves, often in large clusters, to the root system of the cordgrass *Spartina alterniflora* (Bertness 1999). By doing so, the mussels further stabilize the fine sediment of the marsh around the root system, aerate the soil, and provide key nutrients for plant growth (Angelini et al 2016). It has further been recognized that larger, stable salt marsh habitats permit the establishment of a greater variety of terrestrial plants in the high marsh (Bruno 2000). To this end, researchers have explored site-by-site variation in marsh ecosystems of upland plant diversity (Kunza & Pennings 2008). The stability of salt marsh ecosystems is also a measure of coastal protection from storms, productivity for fisheries, and opportunities for tourism (Bertness 1999).

Work has also been done to understand how marine recruitment into these ecosystems may affect the trajectory of a marsh. Robinson et al (2009) used mitochondrial diversity across a well-studied marsh ecosystem in Georgia to explore patterns of recruitment and diversity from a series of mainland, intermediate, and ocean-exposed salt marsh sites. Among the results of this work was a statistical association between the mitochondrial diversity of the ribbed mussel *G. demissa* and the upland plant diversity at a small sample of locations in the Georgia Coastal Ecosystems LTER. This result would suggest that marine larval inputs help drive marsh stability, providing a mechanistic linkage to a community-wide response in biodiversity. Needless to say, that linkage would be important if it is robust, but that result was based on a small spatial sample of *G. demissa* and regional marsh habitats. Here we evaluate this hypothesis from Robinson et al (2009) with a broader sample of marsh habitats in southern Georgia. These sites are near the ‘trailing edge’ of the distributional range of *G. demissa*; it is also likely that these locations will begin to reflect the effects of climate-driven redistribution of marine diversity (Sunday et al 2012), with particular populations or types shifting poleward.

This trailing edge of *G. demissa* is therefore worth evaluating for another reason: three decades ago, Sarver et al (1992) had firmed up the recognition that the southern ribbed mussel, *G. granosissima*, is genomically and phenotypically distinct with likely differences in nutrient deposition and metabolic effects on the sediment around them. The northernmost distribution of *G. granosissima* at that time was close to the southernmost sample of *G. demissa* in the Robinson *et al*. (2009) study and a paired dataset from Díaz-Ferguson et al. (2009). Because few studies have evaluated the presence of *G. granosissima* on the Atlantic coast of Florida (for example, see Walker et al 2019), it is not clear where the current boundary or transition between the two species is. Thus, we also use mitochondrial diversity as a proxy for the presence of *G. granosissima*, which is about 12% divergent from *G. demissa* at the COI gene region (JPW, results not shown) - recognizing that mussels in the family Mytilidae are notable for hybridization at transition zones.

## Methods

Our methods of collection and mitochondrial sequencing follow Robinson *et al*. (2009). Sites were selected to correspond to locations surveyed in Kunza & Pennings (2008), as well as an additional sample near St. Augustine, Florida, to see how diversity may have changed since the 2007 collection from that region represented in Diaz-Ferguson et al (2009). The series of sampled locations is listed in Table 1.

**Table 1.**
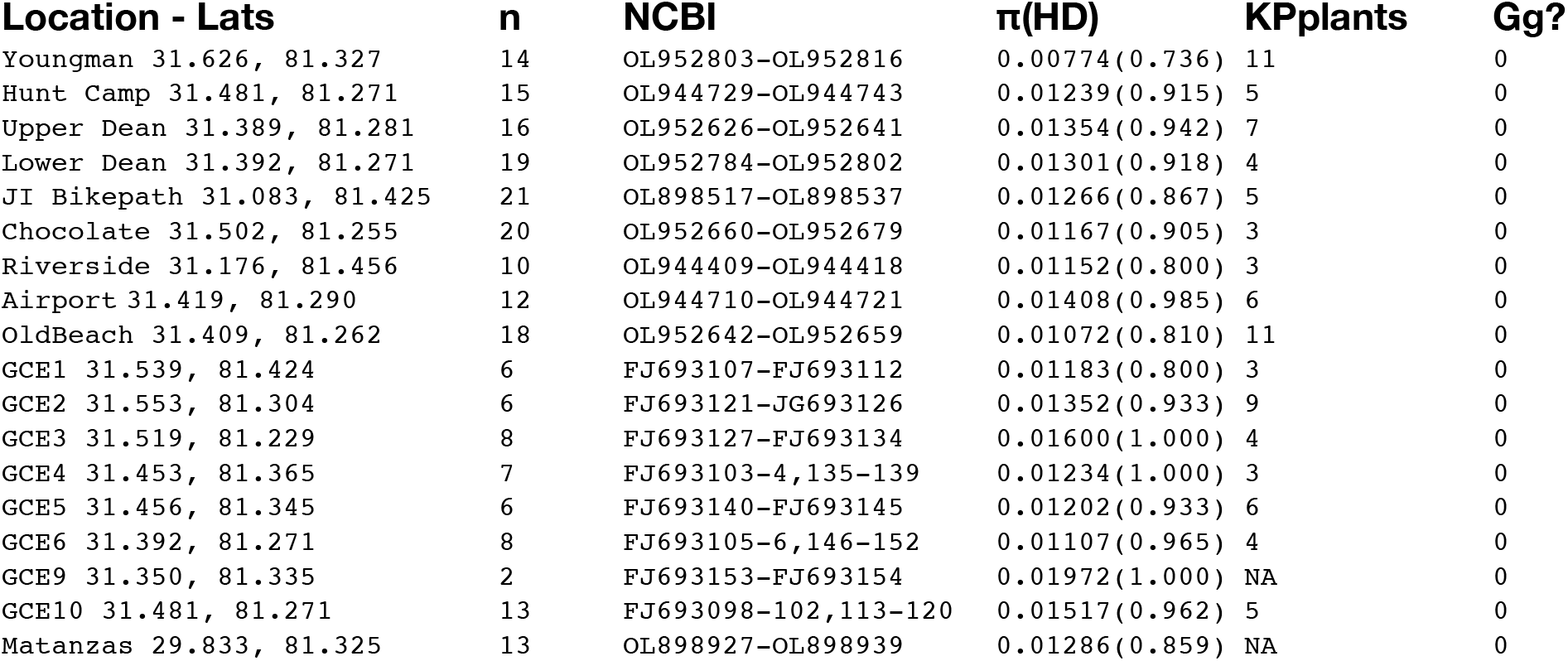
The locations surveyed in this study; sites listed as GCEx are from Robinson et al. (2009), the others were surveyed in 2019-2021. Sample size, NCBI accession numbers, nucleotide and haplotype diversity, the number of plant species identified in Kunza & Pennings (2008) proximal to that site, and presence of G. granosissima are listed for each sample.

| Location - Lats | n | NCBI | $\pi$(HD) | KPplants | Gg? |
| --- | --- | --- | --- | --- | --- |
| Youngman 31.626, 81.327 | 14 | OL952803-OL952816 | 0.00774(0.736) | 11 | 0 |
| Hunt Camp 31.481, 81.271 | 15 | OL944729-OL944743 | 0.01239(0.915) | 5 | 0 |
| Upper Dean 31.389, 81.281 | 16 | OL952626-OL952641 | 0.01354(0.942) | 7 | 0 |
| Lower Dean 31.392, 81.271 | 19 | OL952784-OL952802 | 0.01301(0.918) | 4 | 0 |
| JI Bikepath 31.083, 81.425 | 21 | OL898517-OL898537 | 0.01266(0.867) | 5 | 0 |
| Chocolate 31.502, 81.255 | 20 | OL952660-OL952679 | 0.01167(0.905) | 3 | 0 |
| Riverside 31.176, 81.456 | 10 | OL944409-OL944418 | 0.01152(0.800) | 3 | 0 |
| Airport 31.419, 81.290 | 12 | OL944710-OL944721 | 0.01408(0.985) | 6 | 0 |
| OldBeach 31.409, 81.262 | 18 | OL952642-OL952659 | 0.01072(0.810) | 11 | 0 |
| GCE1 31.539, 81.424 | 6 | FJ693107-FJ693112 | 0.01183(0.800) | 3 | 0 |
| GCE2 31.553, 81.304 | 6 | FJ693121-JG693126 | 0.01352(0.933) | 9 | 0 |
| GCE3 31.519, 81.229 | 8 | FJ693127-FJ693134 | 0.01600(1.000) | 4 | 0 |
| GCE4 31.453, 81.365 | 7 | FJ693103-4,135-139 | 0.01234(1.000) | 3 | 0 |
| GCE5 31.456, 81.345 | 6 | FJ693140-FJ693145 | 0.01202(0.933) | 6 | 0 |
| GCE6 31.392, 81.271 | 8 | FJ693105-6,146-152 | 0.01107(0.965) | 4 | 0 |
| GCE9 31.350, 81.335 | 2 | FJ693153-FJ693154 | 0.01972(1.000) | NA | 0 |
| GCE10 31.481, 81.271 | 13 | FJ693098-102,113-120 | 0.01517(0.962) | 5 | 0 |
| Matanzas 29.833, 81.325 | 13 | OL898927-OL898939 | 0.01286(0.859) | NA | 0 |

To improve the efficiency of PCR amplification, we modified our approach with *Geukensia*-specific primers for the COI gene region. The primers are Geuk-COI-F (CACCCTGGTAACTCTCTTTT) and Geuk-COI-R (ATCACCTCCACCAATAGGAT), using a 50° annealing temperature and protocols otherwise the same as Robinson *et al*. (2009). We isolated DNA from tissue samples using the Puregene protocol, and typically dilute the template to <50ng/μl prior to PCR. Successful reactions were treated with exonuclease and phosphatase prior to submission for DNA sequencing at Psomagen.

Sequence data were evaluated, trimmed, edited and aligned in CodonCode Aligner v9. All data were aligned with representative COI sequence data from *G. granosissima* (NCBI AY621919-AY621922) using the Geneious 11 software to establish possible mitotypes matching the southern congener. Sequences were then analyzed by site in DNAsp v6 (Librado and Rozas, 2009), evaluating nucleotide diversity (π) as a proxy for overall diversity in the system. We also evaluated standard neutrality tests (Tajima’s D), not to explore neutrality but to recognize places where admixture may generate unusual dynamics of spatial diversity, as in Ewers and Wares (2012). We also calculated pairwise divergence, using Kst and Hudson’s Snn, in DNAsp among sample locations.

The results of nucleotide diversity are evaluated against data on high marsh plant diversity from Kunza & Pennings (KP); in Robinson et al (2009), the GCE study sites were compared with the nearest study site of KP (within 3km); the new samples are at the KP data sites. We ask if the hypothesis of correlated diversity as in Robinson et al (2009) holds up with greater sampling from this region using a Pearson’s correlation test performed in R with the package *ggpubr* (Kassambara 2020).

## Results

Across the spatial domain of this study, there is no statistically significant pattern of divergence (Snn 0.057, p = 0.715; Kst −0.019, p = 0.918). Sequence data are archived at NCBI (see Table 1). The correlation between π and high marsh plant diversity is not positive,nor with haplotype diversity and high marsh plant diversity. A Pearson’s correlation test generates a value for t = −1.52 (14 df), p = 0.151, with a negative correlation (−0.376; Fig 1).Very similar results are obtained with haplotype diversity (Table 1). We thus refute the hypothesis of Robinson et al (2009) regarding *Geukensia* diversity and species diversity of the local plant community.

**Figure 1.**
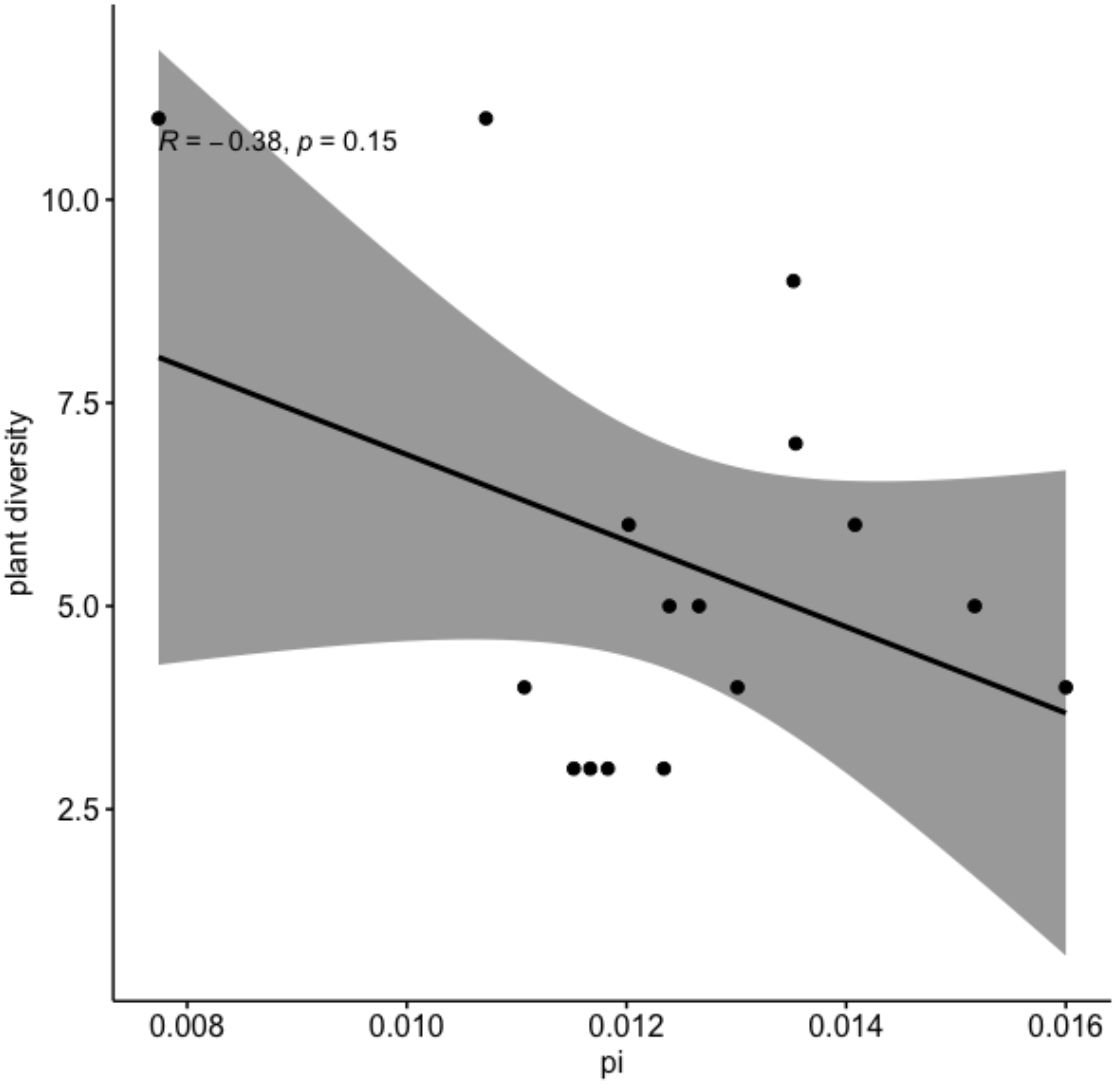
Pearson’s correlation test and distribution of nucleotide diversity in G. demissa and nearby high marsh plant species diversity.

There were zero instances of *G. granosissima* mitotypes throughout these samples, ranging as far south as St. Augustine, Florida (site along west side of Matanzas causeway). We note that the sample from Diaz-Ferguson et al (2009) near St. Augustine, FL also returned no evidence of *G. granosissima*.

## Discussion

It is perhaps unsurprising that the speculative relationship described between mitochondrial diversity in the ribbed mussel and ecosystem-level plant species diversity in Robinson et al (2009) does not hold up to additional data. That result was an outcome of asking a distinct question about larval transport in the marsh ecosystem, but of interest given our need to monitor and protect marsh ecosystems (Bertness 1999). In hindsight, the data in that study for mitochondrial diversity of *G. demissa* were thin enough that extrapolating to such a correlation was not appropriate (*n.b*., J.P.W. was senior author on the Robinson study).

It is, however, surprising that we have so far recovered no evidence of *G. granosissima* in these data. Sarver et al (1992) collected mussels near Ormond Beach, FL (approximately 29.3°N) that included 23 genotypically-identified *G. demissa*, 3 *G. granosissima*, and 6 genotypically-intermediate individuals. Thirty years later, with all we know of climate-driven range shifts in marine organisms, one would expect some evidence of *G. granosissima* to have been picked up accidentally in the marsh surveys of Diaz-Ferguson et al (2009) or the present study in northern Florida. Typical rates of poleward distributional shift have been estimated at >70km/ decade (Poloczanska et al. 2016). Though distributional boundaries do not always track recognizable environmental shifts (Bell et al. 2014, Wares & Skoczen 2019), there is the potential for the distinct diversity of *G. granosissima* to have quietly shifted into marshes – assuming other marine invertebrate shifts as proxies – as far north as Charleston, SC. The current data suggest that this has not happened.

Earlier work in hybrid studies of *Mytilus*, at boundaries between populations in the wild, showed that the mitotypes could move via hybridization - even disrupting typical sex associations with the doubly-uniparental inheritance of mitochondria in mytilids (Rawson et al 1996). However, repeatedly it has been seen to have the mitotypes of one species appear in another in *Mytilus* (Edwards & Skibinski 1987; Kijewski et al 2006, Stuckas et al 2009). Hybrid mussels may be at a fitness disadvantage away from hybrid zones, thus maintaining the genetic integrity of species pairs that overlap (Wood et al 2003). As no additional work on the dynamics of *G. demissa* and *G. granosissima* has been done since the Sarver et al (1992) study, much remains to be known about this pair of mytilid bivalves and their interactions.

A recent attempt to re-establish the transition region with *G. granosissima* by JPW was unsuccessful in locating any mussels that were phenotypically distinct from *G. demissa*, using traits noted in Sarver et al (1992), in the *Spartina*-dominated marshes near the last observed location (Ormond Beach). Considerable development and ‘renewal’ of marsh habitats in this part of Florida may be one disruptive factor. This has long been a concern for potentially disrupting natural biological patterns, as Sumner (1926) noted “[the] orgy of land speculation which prevails at present in Florida, and which promises to upset natural conditions throughout most of the state.” Other recent studies have confirmed the tendency for genomic disruptions – clines and other hybrid zone dynamics – to frequently center on habitat disruption (Wares et al. 2021), possibly pinning those transition zones in place for the time being.

However, it is also possible that the southern species *G. granosissima* is environmentally associated with mangrove habitats to the south of sites examined by J.P.W.; Walker et al (2019) noted ribbed mussels in both habitats of the saltmarsh-mangrove ecotone of this region in northeastern Florida, but did not distinguish between the two congeneric mussels. Numerous studies of *G. granosissima* in the Keys, western Florida, and the Gulf of Mexico note the association with mangroves (Petuch and Myers 2014, p25; Hudson 2017, Honig et al 2014) but it is also found with *Spartina* in the Gulf of Mexico (Rietl et al 2017).

While our data may not immediately change our understanding of salt marsh diversity or interactions, they do act as a baseline of mitochondrial diversity in this region - with data collected in 2007 and 2019-2021 - that will be important for continued analysis of the transition between ecosystems associated with changing climate on the Atlantic southeastern coast of the US. Particularly given the important roles of sediment stabilization and nutrient cycling that ribbed mussels provide to salt marshes, recognizing as other species move in to take that role – particularly if there are some inferred metabolic differences (Sarver et al 1992) – will be of key importance.

## Acknowledgments

This work was supported by a UGA Vice President for Research faculty seed grant in 2020 and the Elaine Lutz Foundation. I. G. F. was supported by the Center for Undergraduate Research Opportunities at UGA. We appreciate the help from B. Freeman, J. Bennett, T. Osborne, C. Angelini, and others in aiding with research or enthusiasm.

